# Multiple genetic changes underlie the evolution of long-tailed forest deer mice

**DOI:** 10.1101/041699

**Authors:** Evan P. Kingsley, Krzysztof M. Kozak, Susanne P. Pfeifer, Dou-Shuan Yang, Hopi E. Hoekstra

## Abstract

Understanding both the role of selection in driving phenotypic change and its underlying genetic basis remain major challenges in evolutionary biology. Here we focus on a classic system of local adaptation in the North American deer mouse, *Peromyscus maniculatus*, which occupies two main habitat types, prairie and forest. Using historical collections we demonstrate that forest-dwelling mice have longer tails than those from non-forested habitats, even when we account for individual and population relatedness. Based on genome-wide SNP capture data, we find that mice from forested habitats in the eastern and western parts of their range form separate clades, suggesting that increased tail length evolved independently from a short-tailed ancestor. Two major changes in skeletal morphology can give rise to longer tails—increased number and increased length of vertebrae—and we find that forest mice in the east and west have both more and longer caudal vertebrae, but not trunk vertebrae, than nearby prairie forms. Using a second-generation intercross between a prairie and forest pair, we show that the number and length of caudal vertebrae are not correlated in this recombinant population, suggesting that variation in these traits is controlled by separate genetic loci. Together, these results demonstrate convergent evolution of the long-tailed forest phenotype through multiple, distinct genetic mechanisms (controlling vertebral length and vertebral number), thus suggesting that these morphological changes—either independently or together—are adaptive.

## INTRODUCTION

Understanding both the ultimate and proximate mechanisms driving adaptation remains a major challenge in biology. One way in which researchers have implicated a role of natural selection in driving phenotypic change is to show the repeated evolution of a trait in similar environments. Such correlations, when controlled for phylogenetic relatedness, can provide evidence for selection rather than stochastic process driving the evolution of a trait of interest (Felsenstein 1985; Harvey & Pagel 1991). Because trait evolution depends on genetic change, we can gain a deeper understanding of adaptation by also studying its underlying genetic basis. For example, what are the genetic mechanisms responsible for phenotypic change, and are they the same or different across independently evolving populations? Understanding the genetic underpinnings of convergent evolution can inform us about the roles of selection and constraint and how these processes may affect evolutionary outcomes (Arendt & Reznick 2008; Manceau *et al*. 2010; Elmer & Meyer 2011; Losos 2011; Martin & Orgogozo 2013; Rosenblum *et al*. 2014). Thus, by studying both the organismal-level processes and the underlying genetic mechanisms driving evolutionary change, we have a much more complete picture of the adaptive process.

Variation among populations of the deer mouse, *Peromyscus maniculatus*, provides a system for understanding the both the genetic and organismal basis of evolution by local adaptation. This species has the widest range of North American mammals (Hall 1981), and populations are adapted to their local environments in many parts of the range (Fig 1A; e.g., Dice 1947; Hammond *et al*. 1988; Storz *et al*. 2007; Linnen *et al*. 2009; Bedford & Hoekstra 2015). Perhaps most strikingly, following the Pleistocene glacial maximum in North America, mice migrated from southern grassland habitat northward and colonized forested habitats, where they have become more arboreal (Hibbard 1968). Natural historians have long recognized two ecotypes of deer mice: forest-dwelling and prairie-dwelling forms (e.g., Dice 1940; Blair 1950). These ecotypes are differentiated both behaviorally and morphologically: mice found in forests tend to have smaller home ranges (Blair 1942; Howard 1949), prefer forest habitat over prairie habitat (Harris 1952), and prefer elevated perches (Horner 1954) as well as have bigger ears, longer hind feet, and longer tails (Osgood 1909; Blair 1950; Horner 1954) than prairie forms. However, it is unknown if the current, widespread forest populations reflect a single origin of the arboreal morphology that has spread across the continent, or if independent lineages have converged on the forest phenotype.

**Figure 1.**
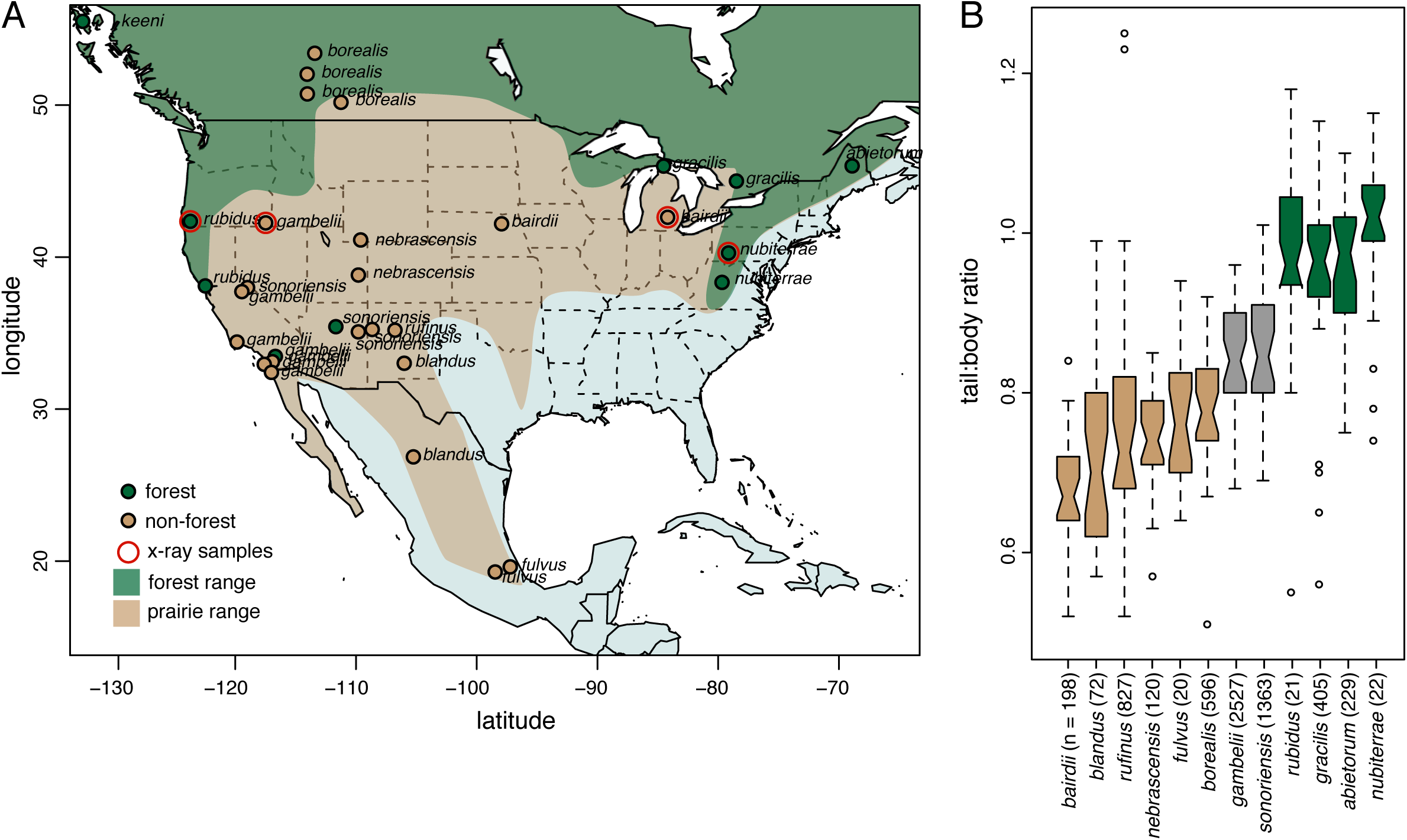
Deer mouse geography and tail length variation. **A**. Map of North America showing the roughly defined range of *P. maniculatus*. Broad-scale forest (green) and prairie (tan) range limits are shown, following Osgood (1909) and Hall (1981). Each dot represents a collecting locale from which one to nine samples were obtained. Dot color represents the local GIS land cover-defined habitat of the site (green or tan). Mice from all sample locales were used in subsequent genetic analyses. The red outline for “x-ray samples” indicates that we used samples from those locations for the comparison of vertebral number and length. **B**. Box-and-whisker plot of tail:body length ratio variation among deer mouse subspecies from museum collections. Box color indicates local habitat, in which samples were collected based on GIS land cover data for those subspecies; tan = prairie, grey = mixed, green = forest.

Of the morphological traits associated with forest habitats, arguably the best recognized is tail length, and evidence suggests that increased tail length is an adaptive response to increased arboreality. Previous work has shown that deer mice use their tails extensively while climbing. Horner (1954) carried out elegant experiments on climbing behavior in *Peromyscus*, and not only found a correlation between tail length and climbing ability, but also provided experimental evidence that within *P. maniculatus*, forest mice are more proficient climbers than their short-tailed counterparts and rely on their tails for this ability. Moreover, in two other *Peromyscus* species, tail length correlates with degree of arboreality (Smartt & Lemen 1980), and climbing ability has been shown to be heritable (Thompson 1990).

Here, we investigate the evolution of the deer mouse tail in several complementary ways. First, we reconstruct phylogeographic relationships among 31 populations of *P. maniculatus* to test hypotheses about the evolution of tail length. We show that forest-dwelling deer mice do not belong to a single phyletic group or genetic cluster, and thus that long-tailed forest forms appear to have independently converged. We also demonstrate that the evolution of longer tails is correlated with living in forest habitats, even when taking account of shared history and gene flow among populations. Second, we investigate the morphological basis of tail length differences in two geographically distant populations, implicating similar but quantitatively different mechanisms for the generation of distinct tail morphologies in eastern and western forest populations. Finally, we show that differences in the constituent traits of tail length between forest and prairie mice, despite correlation of these traits in the wild, are genetically separable in recombinant laboratory populations. Together, these results strongly suggest that natural selection maintains multiple locally adapted forest populations.

## METHODS

### Samples of museum specimens

To quantify the degree of variation in overall tail length in this widespread species, records of *P. maniculatus* were downloaded from the Mammal Network Information System (MaNIS 2015), the Arctos database (Arctos 2015), and individual museum databases. Because nearly all specimens were present in the collection as prepared skins, we considered the original collector’s field data to be the most reliable source of measurements. We excluded all specimens labeled as “juvenile”, “subadult” or “young adult” or those having any tail abnormalities or injuries. We also removed any individuals with total length below 106 mm or tail length below 46 mm, which are considered the adult minima for this species (Hall 1981; Zheng *et al*. 2003). In total, we gathered data from 6,400 specimens.

To assign these specimen to subspecies, we scanned the original distribution maps from Hall (1981) and georeferenced them in ArcGIS v. 9.2 (ESRI 2007). Subspecies were assigned based on original identification when available and on mapping the coordinates digitally. *Peromyscus maniculatus* has highly stable subspecies ranges, and subspecific identity can be determined reliably based on appearance and location (King 1968; Gunn & Greenbaum 1986; Hall 1981).

### Morphometric measurements and statistics

We calculated two types of dependent variables. First, we calculated the ratio of tail length to body length for all individuals. Second, we addressed potential non-linear scaling of the two measures by fitting a linear model of tail length *vs*. body length (Fox & Weisberg 2011). Models, including log, square, and Box-Cox transformations of the response variable, as well as quadratic and cubic terms for the predictor, were compared using ANOVA and the adjusted R^2^ values. Because the best model had very low explanatory power (R^2^ = 0.11), we decided that simple ratios are the best statistic for this dataset. Because the ratios were not normally distributed (p < 0.001, Kolmogorov-Smirnoff test), the means of plains, forest, and unclassified forms were compared with the Kruskal-Wallis non-parametric ANOVA. The R package *car* was used for all computation (R Development Core Team 2005; Fox & Weisberg 2011).

### Morphology and habitat designation

Because skeletal preparations of museum specimen most often don’t have complete tails (e.g., the smallest vertebrae are often lost during sample preparation), we conducted more detailed analyses on whole mouse specimens (i.e., frozen or fluid-preserved specimens). We focused the next set of genomic and morphometric analyses on 80 individuals from our lab collection as well as loans from researchers and museums, covering multiple regions included in the above analyses (see Supp. Table 1 for details). The sampling encompassed the range of *P. maniculatus* and included 31 locations, spread across multiple habitats (Fig. 1A).

We assigned these animals to local habitat types based on their specific sampling locations: we used ArcGIS (ESRI) to extract land cover (i.e., habitat type) information from the North American Land Change Monitoring System 2010 Land Cover Database (NALCMS 2010) with a 1 km-radius buffer around each sampling location. We split land cover categories into forest and non-forest/prairie designations for all analyses: we called classes 1-6 and 14 forest (Temperate or sub-polar needleleaf forest, Sub-polar taiga needleleaf forest, Tropical or sub-tropical broadleaf evergreen forest, Tropical or subtropical broadleaf deciduous forest, Temperate or sub-polar broadleaf deciduous forest, Mixed Forest, Wetland) and others non-forest/prairie (Tropical or sub-tropical shrubland, Temperate or sub-polar shrubland, Tropical or sub-tropical grassland, Temperate or subpolar grassland, Cropland, Urban and Built-up).

### Array-based capture and sequencing of short-read libraries

To assess genome-wide population structure, we used an array-based capture library and sequenced region-enriched genomic libraries using an Illumina platform (Gnirke *et al*. 2009). Our MYbaits (MYcroarray; Ann Arbor, MI) capture library sequences include probes for 5114 regions of the *Peromyscus maniculatus* genome, each averaging 1.5kb in length (totaling 5.2Mb of unique, non-repetitive sequence; see Domingues *et al*. [2012] and Linnen *et al*. [2013]). We extracted genomic DNA using DNeasy kits (Qiagen; Germantown, MD) or the Autogenprep 965 (Autogen; Holliston, MA) and quantified it using Quant-it (Life Technologies). 1.5μg of each sample was sonically sheared by Covaris (Woburn, MA) to an average size of 200bp, and Illumina sequencing libraries were prepared and enriched following Domingues *et al*. (2012). Briefly, we prepared multiplexed sequencing libraries in five pools of 16 individuals each using a “with-bead” protocol (Fisher *et al*. 2011) and enriched the libraries following the MYbaits protocol. We pulled down enrichment targets with magnetic beads (Dynabeads, Life Technologies; Carlsbad, CA), PCR amplified with universal primers (Gnirke *et al*. 2009), and generated 150bp paired end reads on a HiSeq2000 (Illumina Inc.; San Diego, CA).

### Sequence alignment

We preprocessed raw reads (fastq files) by removing any potential non-target species sequence (e.g., from sequence adapters) and by trimming low quality ends using cutadapt v. 1.8 (Martin 2011) and Trim Galore! v. 0.3.7 (www.bioinformatics.babraham.ac.uk/projects/trim_galore) before aligning them to the Pman_1.0 reference assembly (www.ncbi.nlm.nih.gov/assembly/GCF_000500345.1) using Stampy v. 1.0.22 (Lunter & Goodson 2011). We removed optical duplicates using SAMtools v. 1.2 (Li *et al*. 2009), retaining the read pair with the highest mapping quality. Prior to calling variants, we performed a multiple sequence alignment using the Genome Analysis Toolkit v. 3.3 (GATK) IndelRealigner tool (McKenna *et al*. 2010; DePristo *et al*. 2011; Van der Auwera *et al*. 2013). Following GATK’s Best Practice recommendations, we computed Per-Base Alignment Qualities (BAQ) (Li 2011), merged reads originating from a single sample across different lanes, and removed PCR duplicates using SAMtools v. 1.2 (Li *et al*. 2009). To obtain consistent alignments across all lanes within a sample, we conducted a second multiple sequence alignment and recalculated BAQ scores. We finally limited our dataset to proper pairs using SAMtools v. 1.2.

### Variant calling and filtering

We used GATK’s HaplotypeCaller to generate initial variant calls via local *de novo*1 assembly of haplotypes. We combined the resulting gvcf files using GATK’s CombineGVCFs command and jointly genotyped the samples using GATK’s GenotypeGVCFs tool. Post genotyping, we filtered these initial variant calls using GATK’s VariantFiltration in order to minimize the number of false positives in the dataset. In particular, we applied the following set of filter criteria: we excluded SNPs for which three or more variants were found within a 10bp surrounding window (clusterWindowSize=10); there was evidence of a strand bias as estimated by Fisher’s exact test (FS>60.0); the read mapping quality was low (MQ<60); the mean of all genotype qualities was low (GQ_MEAN<20). We limited the variant dataset to biallelic sites using VCFtools (Danecek *et al*. 2011) and excluded genomic positions that fell within repeat regions of the reference assembly (as determined by RepeatMasker [Smit *et al*. 2013–2015]). We excluded genotypes with a genotype quality score of less than 20 (corresponding to P[error] = 0.01) or a read depth of less than four using VCFtools to minimize genotyping errors. In addition, we filtered variants on the basis of Hardy Weinberg Equilibrium (HWE): a p-value for HWE was calculated for each variant using VCFtools, and variants with p < 0.01 were removed.

### Genetic principal components analysis (PCA)

To assess genetic structure across the range of *P. maniculatus*, we first used SMARTPCA and TWSTATS (Patterson *et al*. 2006) with the genome-wide SNP data (after filtering for variants genotyped in >50% of individuals, the final dataset contained 7,396 variants) from the enriched short-read libraries described above. We detected significant structure in the first two principal components (p < 0.05 by Tracy-Widom theory, as applied in TWSTATS). To further visualize genetic relationships inferred by the PCA, we generated a neighbor-joining tree by computing Euclidean distances between individuals in the significant eigenvectors of the SMARTPCA output. The Python code to produce these trees can be found at *github.com/kingsleyevan/phylo_epk*.

### Phylogeography and monophyly of forest forms

Relationships and gene flow between the populations of P. *maniculatus* were further investigated using the Ancestral Recombination Graph approach in the software TreeMix v. 1.12 (Pickrell & Pritchard 2012). As phylogenetic inference requires quartets, we filtered the capture data to include only the variable sites with high quality data for at least four individuals (14,076 SNPs) and converted it using PLINK v. 1.8 (Purcell *et al*. 2007). We inferred a bifurcating tree from the allele frequency spectrum. To account for linkage disequilibrium, we grouped SNPs into blocks of 10 or 100; block size did not affect the results. We subsequently fitted 10 or 20 migration events between *a priori* designated populations and estimated the magnitude and significance of admixture by jackknifing. Command line: *treemix-i P.maniculatus.tmix.gz-o P.maniculatus.k100.m10-root leucopus-k 100-m 10-noss-se*.

The VCF was converted to a fasta alignment using PGDSpider v. 2.0 (Lischer & Excoffier 2012). We generated a phylogenetic tree in RAxML v. 8 under the GTR+G model with 500 bootstrap replicates and a correction for ascertainment bias (Stamatakis 2014). We then tested the hypothesis that long-tailed forest individuals are monophyletic, either across the continent or within each of the western and eastern clades, by constraining the tree inference and comparing the likelihood using the Approximately Unbiased test (AU) (Shimodaira 2002) in CONSEL v. 0.2 (Shimodaira 2001).

### Mitochondrial DNA analysis

Mitochondrial sequences were generated for a set of 106 samples, including 35 of those used in the capture experiment as well as *P. polionotus, P. keeni and P. leucopus* as outgroups (Supp. Table 2). Amplification and sequencing were performed following Hoekstra *et al*. (2005). Briefly, we amplified the region spanning the single-exon gene CO3 and ND3, with the intervening Glycine tRNA, in a single PCR, and Sanger-sequenced using the primers 5’ CATAATCTAATGAGTCGAAATC 3’ (forward) and 5’ GCWGTMGCMWTWATYCAWGC 3’ (reverse). We aligned the resultant 1190 bp sequences using MUSCLE (Edgar 2004). The ML tree estimation and monophyly testing were carried out as above, with a separate partition defined for each codon position.

### Within-species comparative analysis

To test for a correlation between tail length and habitat type, we used a comparative approach that accounts for the phylogenetic relationships among samples. First, we calculated standard Phylogenetically Independent Contrasts (Felsenstein 1985), based on both the nuclear and mitochondrial ML phylogenies, using the R package GEIGER (Harmon 2008). However, because we are comparing populations within a single species, this analysis is potentially confounded by non-independence because of gene flow in addition to shared ancestry (Stone *et al*. 2011). Nonetheless, these results support the conclusions of the two controlled tests described below.

To control for the non-independence of populations outside the context of bifurcating tree, we used a generalized linear mixed model approach as described by Stone *et al*. (2011). We used two measures of genetic similarity. First, we pared our data down to populations from which we had three or more individuals (now six populations) and estimated global mean and weighted F_ST_ between populations using the method of Weir and Cockerham (1984) as implemented in VCFtools (Danecek *et al*. 2011; see “in population contrasts” in Supp. Table 1). Second, we used a measure of genetic similarity (“–relatedness2” in VCFtools: the kinship coefficient of Manichaikul *et al*. 2010) between all individuals in the dataset for which we have tail and body measurements (” in individual contrasts” in Supp. Table 1). For both the population-and individual-level analyses, we used a generalized linear mixed model approach, as implemented in the R package *MCMCglmm* (Hadfield 2010) to test for an effect of habitat (forest vs. non-forest) on population average tail:body ratio or individual tail:body ratio when including F_ST_ or kinship coefficient as a random effect.

### Vertebral morphometrics of wild-caught specimens

To measure tail morphology, we x-ray imaged the skeletons of wild-caught mice from four populations (Fig. 1A [red circles], and see Supp. Table 1 for specimen details). We used a Varian x-ray source and digital imaging panel in the Museum of Comparative Zoology Digital Imaging Facility. Then we used ImageJ’s segmented line tool to measure the lengths of each individual vertebra, starting from the first caudal vertebra and proceeding posterior to the end of the tail. The boundary between the sacral and caudal segments of the vertebral column is not always clear, so for consistency, we called the first six vertebrae (starting with the first sacral-attached) the sacral vertebrae; the caudal vertebrae are all the vertebrae posterior to the sixth sacral vertebra. This approach allowed us to reliably compare vertebrae across individuals.

Because tail length scales with body size in our sample, we fitted a linear model with the *lm* function in R (R Development Core Team 2005) to adjust all vertebral length measurements for body size. We regressed total tail length (R^2^ = 0.62) and the lengths of individual vertebrae (R^2^ = 0.14–0.59) on the sum length of the six sacral vertebrae and used the residuals from the linear fit for all subsequent analyses. We obtained similar results when we regress vertebral length measurements on femur length instead of sacral vertebral length (results not shown).

### F2 intercross trait correlation analysis

To examine the genetic architecture of tail traits, we conducted a genetic cross between a prairie and forest population. All mice were housed in the Hoekstra lab colony at Harvard University. We originally obtained prairie deer mice, *P. m. bairdii*, from the Peromyscus Genetic Stock Center (University of South Carolina), which we crossed with forest deer mice, *P. m. nubiterrae*, that we originally captured in Westmoreland County, Pennsylvania. We mated a single male and female of each subspecies—two mating pairs, one in each cross direction—and used their offspring to establish 10 F1 sibling mating pairs. We then x-ray imaged the resulting 96 F2 offspring and measured their tail morphology as described above (section on Vertebral morphometrics). We used the *lm* function in R to assess correlations in the resulting measurements. All animals were adults between 80 and 100 days old when x-rayed.

## RESULTS

### Forest deer mice do not form a single clade

We examined museum records of 6,400 specimens belonging to 12 subspecies of *Peromyscus maniculatus* from across North America. Nearly all of the forest forms have substantially longer tails than the prairie populations (Kruskal-Wallis test, p = 0.001), often equaling the length of their body (Fig. 1B). We used an array-based capture approach to resequence and call SNPs in >5000 genomic regions in a continent-wide sample of animals from 31 *P. maniculatus* populations, *P. (maniculatus) keeni*, and an outgroup *P. leucopus* (Fig. 1A). We identified 14,076 high-quality SNPs, corresponding to an average of one variant every 1,342.6bp of the reference genome with 24.73% of the genotypes missing. We explored genetic similarity among the sampled individuals by genetic principal components analysis (PCA; Patterson *et al*. 2006) and Maximum Likelihood phylogenetic inference (Fig. 2).

**Figure 2.**
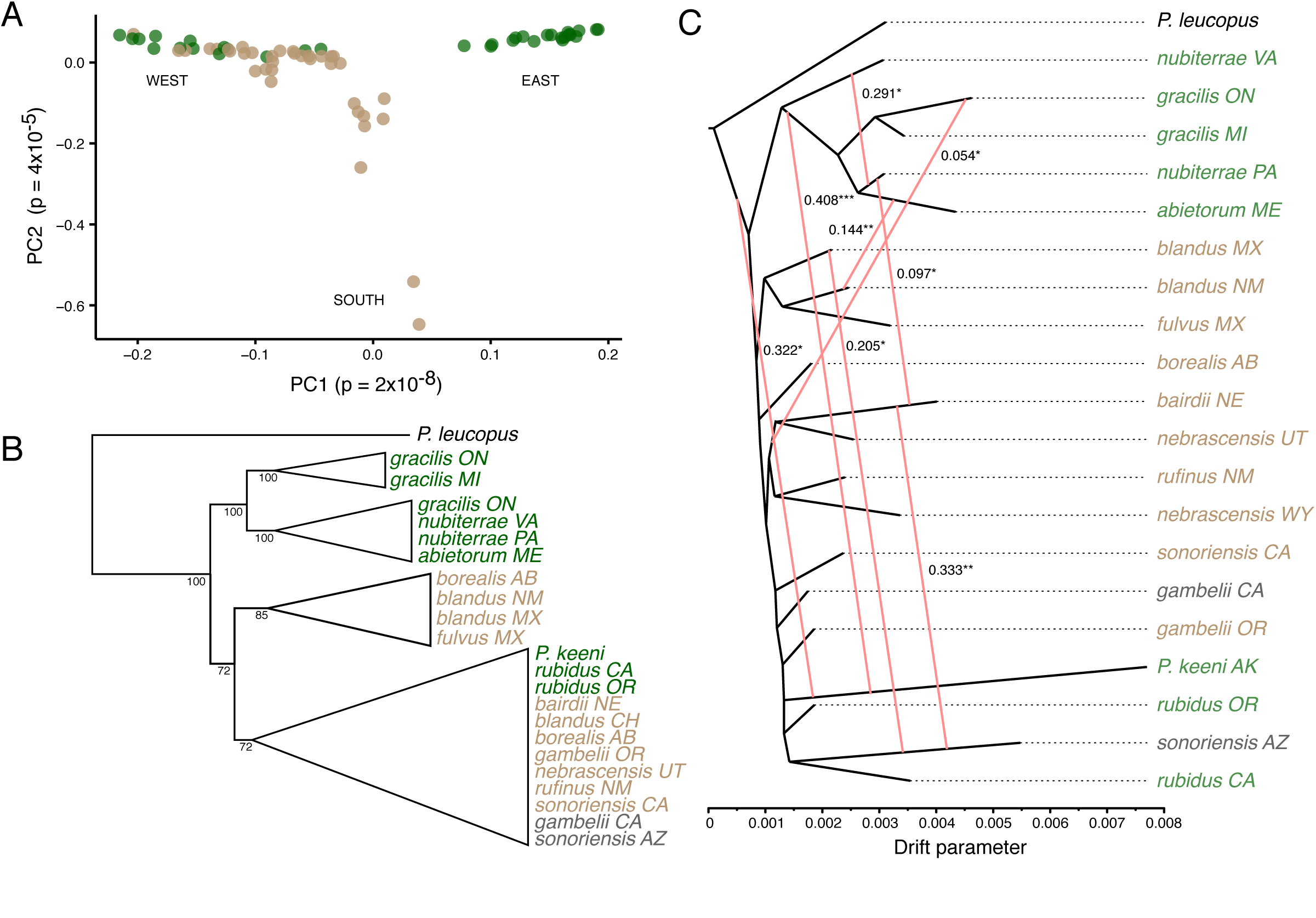
Deer mouse population structure and gene flow. **A**. Plot of the first two Principal Components calculated from a genome-wide sample of 7,396 high-quality SNPs from populations ranging across the continent (sampling locales shown in Fig. 1A). Each dot (n = 80) represents a single individual. **B**. Cladogram based on nuclear Maximum Likelihood phylogeny collapsed to high-confidence clades. Node labels represent bootstrap support. **C**. Phylogenetic tree with mixture events as inferred by TreeMix. Branch lengths correspond to the estimated amount of genetic drift. Numbers indicate the inferred proportion of shared polymorphisms between two lineages connected by a red line. Asterisks indicate gene flow estimates significantly more than zero (*p < 0.05; **p < 0.01; ***p < 0.001). In all panels, colors indicate local GIS land cover-defined habitat (tan = prairie, green = forest, grey = mixed).

Both the PCA and phylogenetic methods show that individuals from forests (as determined by GIS) do not compose a single, monophyletic group. Instead, we see the mice of the putatively derived forest forms clustering with nearby non-forest forms in the PCA (Fig. 2A). Trees based on PCA distances (Supp. Fig. 1), estimated from DNA sequences (both genome-wide SNPs [Fig. 2B; Supp. Fig. 2] and mtDNA [Supp. Fig. 3]), and estimated from allele frequencies (Fig. 2C) all identify multiple origins of forest forms. Monophyly is rejected in ML tests using the nuclear and mitochondrial data (AU test, p<0.001). These results show that forest forms are evolving independently in eastern and western parts of the species range.

**Figure 3.**
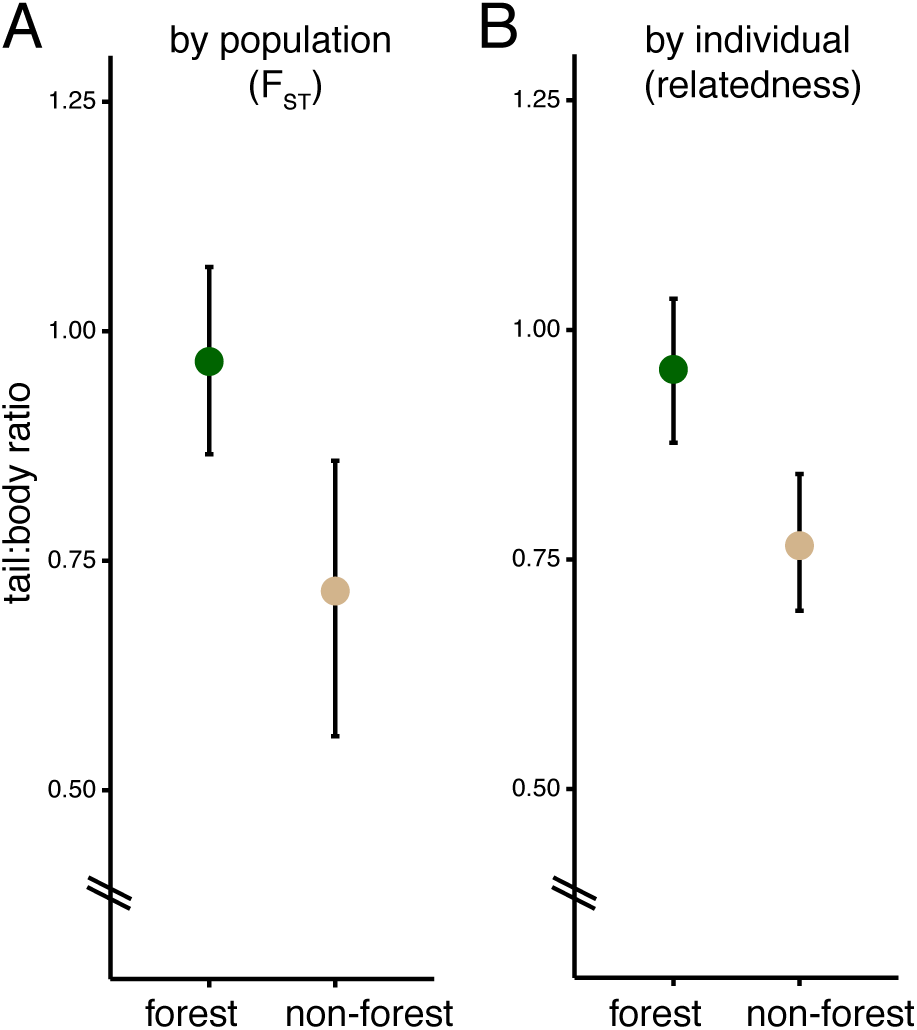
Tail length differences between habitats when accounting for genetic non-independence. **A**. Tail:body length ratios for each population predicted by a generalized linear mixed model taking genetic differentiation (F_ST_) between populations into account. **B**. Tail:body length ratios for prairie and forest individuals predicted by a similar model as in A, but with pairwise genetic relatedness (kinship coefficient [Manichaikul *et al*. 2010]) between individuals taken into account. Error bars represent 95% confidence intervals.

Relationships between populations were also estimated by reconstructing an ancestral recombination graph from allele frequency spectra in TreeMix v. 1.12. As above, four major groups were identified (Mexico, East Coast, Great Plains/North, and West Coast). P. keeni is nested within *P. maniculatus*, although with elevated genetic divergence (Fig. 2C). Moreover, we used these data to estimate connectivity among the populations (i.e., the pairwise proportion of shared ancestral polymorphism). These analyses showed some evidence for gene flow among populations, even across large geographic distances. While most substantial gene flow occurred within a habitat type, in a few cases, we detected significant gene flow between forest and prairie populations, supporting the idea that populations can retain different phenotypes, even in the face of gene flow (Fig. 2).

### Variation in tail length correlates with habitat

To assess whether differences in the length of the tail are significant even when accounting for evolutionary relationships among populations (created by gene flow and/or shared ancestry), we included measures of genetic similarity among populations and individuals in a set of generalized linear mixed models. In these models, we ask whether animals in different habitats have significantly different tail:body ratios when including measures of genetic similarity as random effects.

First, we considered whether forest and prairie populations differ in their mean tail lengths. When taking pairwise F_ST_ between populations into account (Supp. Fig. 4), we find that habitat has a significant effect in our mixed model (p = 0.002 for weighted mean F_ST_, p < 0.001 for global mean F_ST_; fit by Markov Chain Monte Carlo [Hadfield 2010]). Next, we assessed whether individuals from forest and prairie habitats differ in tail length when accounting for genetic similarity. We find a significant effect of habitat on tail:body ratio in our mixed model with kinship coefficient (Manichaikul *et al*. 2010) as a random effect (MCMC; p < 0.001). We show model-predicted population means in Figure 3A and the predicted individual means by habitat in Figure 3B. In addition, phylogenetically independent contrasts show a correlation between habitat type and tail:body length ratio (adjusted R^2^=0.303, p<0.0001). Together, these results robustly show that, even when taking non-independence of populations into account, deer mice from forested habitats do indeed have significantly longer tails than those from prairie habitats.

**Figure 4.**
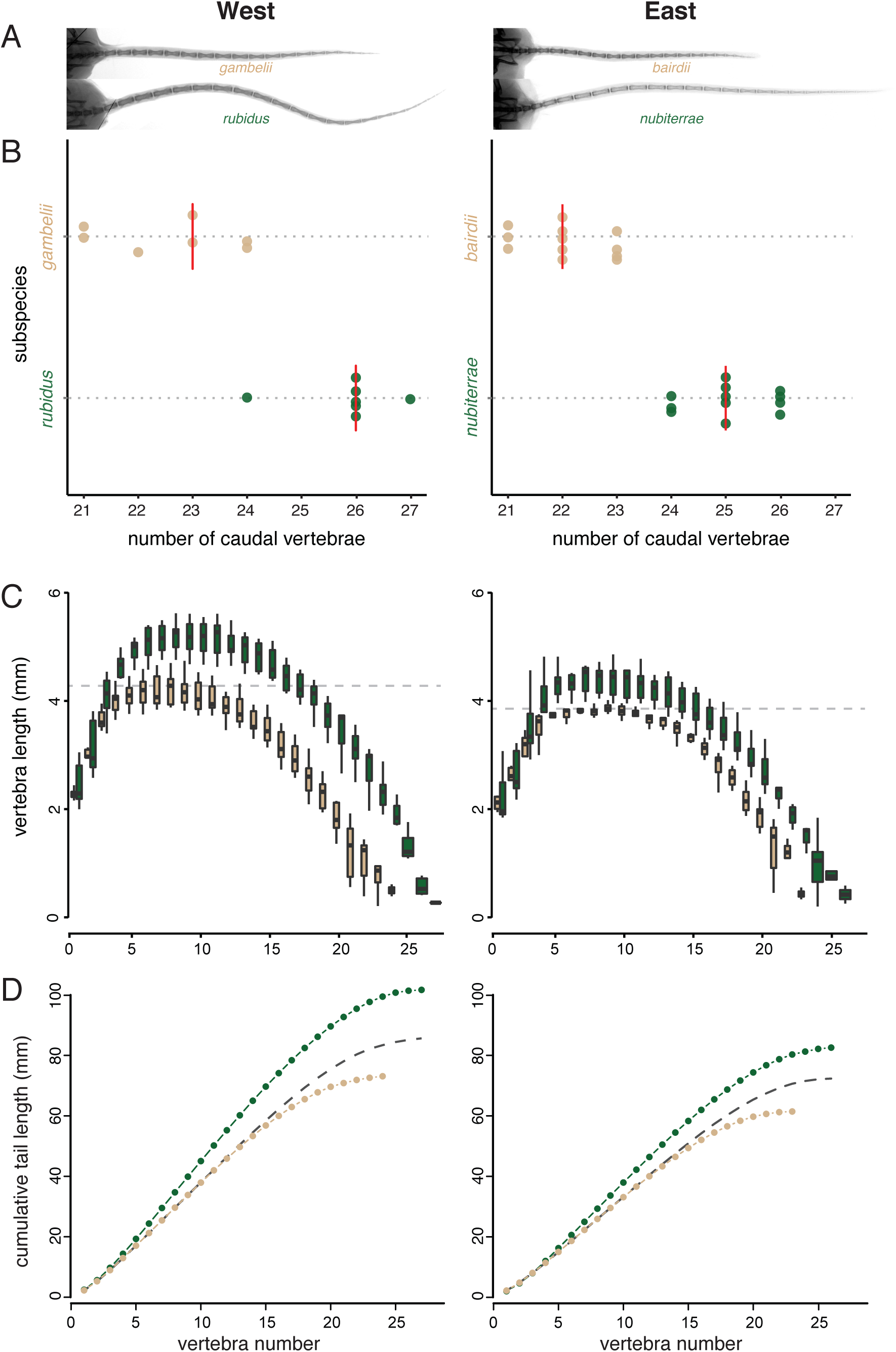
Convergent tail vertebral morphology in eastern and western forest-prairie population pairs. **A**. Representative radiographs of deer mouse tails from the eastern and western population pairs. **B**. Forest mice have more caudal vertebrae than prairie mice in the east and west. Vertical red lines represent medians. Note truncated axis. **C**. Forest mice have longer vertebrae than prairie mice in the east and west. Dashed line represents the median length of the longest prairie vertebra; the segment of the forest tail with longer vertebrae is 4–16 and 4–15 for western and eastern populations, respectively. **D**. Summary of tail vertebral differences between forest and prairie mice. Dots represent mean cumulative tail lengths. Dashed line is the cumulative length of the mean prairie tail with three extra vertebrae added, which represents an estimate of the maximum contribution of difference in number of vertebrae to the difference in total length.

### Convergence in skeletal morphology

We x-rayed specimens from two pairs of geographically distant forest-prairie populations (Fig. 1A [red circles] and “x-ray samples” in Supp. Table 1) and measured their vertebrae. We focused on these four populations because individuals from the two forest populations represent independently evolving forest lineages. Though both forest populations have longer tails than non-forest populations, eastern and western forest mice do not have identical tail lengths: western mice have longer tails, both absolutely and relative to their body size, than eastern forest mice (Fig. 4A-D).

In addition to the total tail length, we measured the number of caudal vertebrae and the lengths of each caudal vertebra. We found that, in both east and west paired comparisons, forest mice had significantly longer tails, significantly more vertebrae, and significantly longer vertebrae (here we tested the single longest vertebrae) than their nearby prairie form (Fig. 4; Wilcoxon tests, p < 1.6e-4 for all comparisons). Notably, we performed all these tests on body-size-corrected data, which means that these forest-prairie differences are not driven by overall differences in body length. We also found significant effects of habitat and subspecies on tail length and longest vertebra length in a two-way analysis of covariance (ANCOVA) on log-transformed data with sacral length as a covariate (p < 6.6e-8 for all effects).

First, we counted the number of caudal vertebrae and found that forest forms have significantly more vertebrae than the prairie forms (Kruskal-Wallis test, X^2^ = 29.0, p = 7.2e-8). In both cases, we found that forest mice had, on average, three more vertebrae than the nearby prairie form. However, the western mice had, on average, one more caudal vertebrae than the eastern population from the same habitat type (Fig. 4B). Importantly, none of the populations can be distinguished by the number of trunk vertebrae (all samples had 18 or 19), showing that vertebral differences are specific to the tail.

We next compared the lengths of the individual caudal vertebrae along the tail and found that many but not all caudal vertebrae are longer in the forest than in prairie mice. In the eastern population pair, caudal vertebrae 4 through 15 had longer median lengths in the forest mice than the longest caudal vertebra in prairie mice. The corresponding segment in the western population pair is caudal vertebrae 4 through 16 (Fig. 4C).

We also estimated the relative contributions of differences in vertebral length and vertebral number to the overall difference in tail length. To do this, we modeled the forest and prairie mice having an equal number of vertebrae by inserting three long vertebrae into the center of the prairie tails. These simulated “prairie+3” tails compensated for 42% and 53% of the average difference in overall tail length between the eastern and western forms, respectively (Fig. 4D). These estimates represent an upper bound of the contribution of the difference in vertebral number relative to vertebral length in these populations. Finally, a linear model of the form *total length ˜ longest vertebra length + number of caudal vertebrae* has an R^2^= 0.98, suggesting that differences in length and number of vertebrae explain nearly all of the difference in total tail length between these populations. Thus, these two morphological traits—length and number of vertebrate—contribute approximately equally to the difference in overall tail length in the western and eastern clades.

### Vertebrae length and vertebrae number are genetically separable

Among the four populations examined above, both forest populations have both more caudal vertebrae and longer caudal vertebrae. Could these differences be correlated in natural populations because they are under the control of the same genetic loci? To test this hypothesis, we generated 96 F2 recombinant individuals from a reciprocal intercross between the eastern pair, long-tailed, forest *P. m. nubiterrae* and short-tailed, prairie *P. m. bairdii*, and generated x-ray images from each second-generation hybrid. Using vertebra measurements from these images, we tested for a correlation between the number of caudal vertebrae and the length of the longest caudal vertebra. We detect no significant correlation (Fig. 5; t = 0.87, df = 94, p = 0.39) between vertebral length and vertebral number in the tails of our F2 animals. A sample of 96 individuals is well powered, allowing an 80% probability of detecting a correlation of r > 0.25 at p < 0.05. These data suggest that these two phenotypic traits—vertebral number and length—are under independent genetic control.

**Figure 5.**
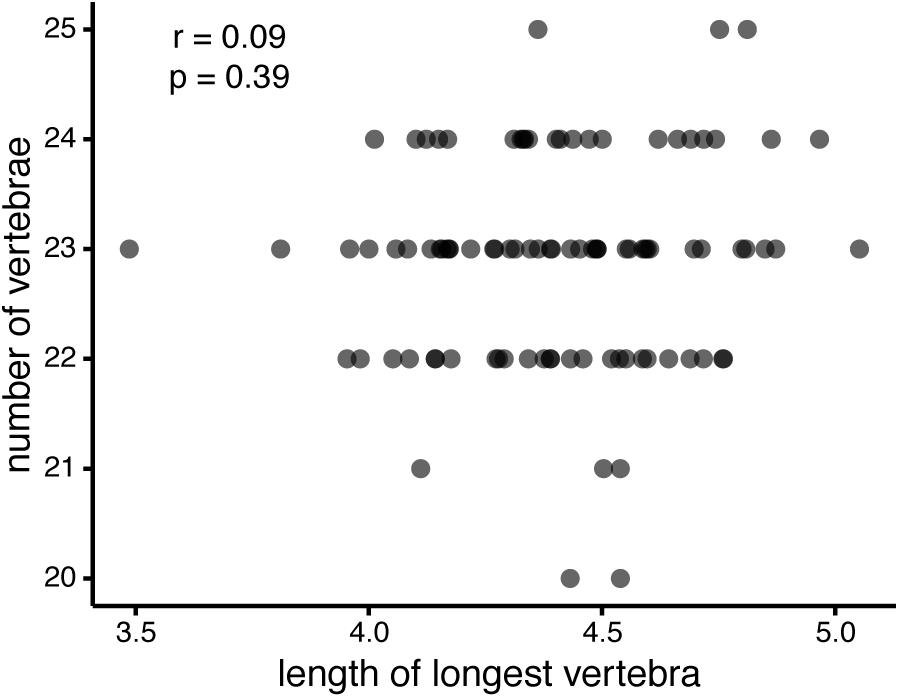
No significant correlation between number and length of caudal vertebrae in a laboratory F2 intercross. Each point is represents the length of the longest caudal vertebra and the number of caudal vertebrae measured from a radiograph of an F2 *nubiterrae* x *bairdii* individual. Ninety-six individuals allows 80% power to detect a correlation of r > 0.25.

## DISCUSSION

In this study, we explored the evolution of a repeated phenotype within a single species. We found that long-tailed forest-dwelling deer mice are evolving independently in eastern and western parts of its range, and that tail length does indeed differ between forest and non-forest habitats, despite gene flow among ecotypes. Furthermore, we showed that longer tails, in both eastern and western forest mice, are a result of differences in both the number of caudal vertebrae and also the lengths of those vertebrae. Finally, we empirically demonstrated that, despite the observation that caudal vertebrae number and length are correlated in nature, the genetic mechanisms producing those differences can be decoupled in a laboratory intercross. Together, these results imply that natural selection is driving differences in tail length, and suggest that proportional changes in both number and length may influence the biomechanics of tail use.

In his large-scale 1950 survey of adaptive variation in *Peromyscus*, W. Frank Blair described two main forms of *P. maniculatus*, one grassland and one forest, supporting previous suggestions (Osgood 1909). Therefore, we examined the phylogeography of this deer mouse species in the context of intraspecific ecological adaptation, and specifically in the framework of the classic prairie-forest dichotomy that has been recognized in this species for over one hundred years. Previous work on the phylogenetic relationships among deer mouse subspecies has suggested convergence in these forms—allozyme (Avise *et al*. 1979) and mitochondrial studies (Lansman *et al*. 1983; Dragoo *et al*. 2006) hinted at a split between eastern and western forest populations—yet none have explicitly considered morphological and ecological context in a continent-wide sampling. Additionally, mitochondrial-nuclear discordance occurs in this species (e.g., Yang & Kenagy 2009; Taylor & Hoffman 2012), which complicates the interpretation of mitochondrial studies. Here, we used > 7000 genome-wide SNPs to directly test for a correlation between habitat and morphology. We chose to use both tree-based and non-tree-based methods, given the difficulty of constructing bifurcating phylogenetic relationships among intraspecific samples from populations experiencing gene flow.

The results of our genetic PCA provide strong evidence, concordant with those of previous studies, that forest mice do not form a single genetic cluster (Fig. 2A). The pattern of genomic differentiation we see in the PCA is roughly similar to that recovered by Avise *et al*. (1979), with a group of populations from eastern North America that are clearly separated from populations in the western half of the continent. We see similar results from maximum likelihood phylogenetic reconstruction among individuals and from allele-frequency-based population trees (Fig. 2B-C). We also infer that there has been gene flow between *P. maniculatus* lineages in our sample. Despite some evidence for admixture, there remains a strong signal of genetic differentiation among populations, and importantly, forest and non-forest populations retain different tail morphologies, suggesting that natural selection maintains these locally adapted phenotypes.

Indeed, when we assigned habitat values to those populations using GIS land cover data, we found that populations captured in forest habitats have longer tails than those in non-forest habitats, even when taking gene flow among populations into account. Thus, we support the hypothesis of Lansman *et al*. (1983): “It is thus very probable that the currently recognized forest-grassland division of *P. maniculatus* does not reflect a fundamental phylogenetic split. Rather, it is more likely that environmental selection pressures have led to the independent evolutionary appearance of these two morphs in different maniculatus lineages.” Our data support at least two, one eastern and one western, independently evolving groups of forest *P. maniculatus*—western North American populations are less differentiated from each other than they are from those in the east, making it difficult to confirm more than a single group of forest mice in the west.

Several lines of evidence suggest that these independent extant forest forms derive from a short-tailed prairie-dwelling ancestor. First, although our SNP-based phylogeny does not imply an obvious ancestral state, our mtDNA phylogeny contains a basal branching prairie lineage (Supp. Fig. 3), consistent with a prairie ancestor to the *P. maniculatus* group. Second, two studies showed that *P. melanotis* is the outgroup to the *P. maniculatus* species group (Bradley *et al*. 2007, Gering *et al*. 2009). This outgroup species, P. melanotis, is found primarily in grassy plains and is characterized by a very short tail (Álvarez-Castañeda 2005). Third, several fossils attributed to *P. maniculatus* and dated to the Pleistocene late-glacial period—during which the *P. maniculatus* group is thought to have radiated—were found in regions likely to be mainly scrub grasslands (Hibbard 1968; Bryant & Holloway 1985). While it is always challenging to infer ancestral phenotypic states, these data suggest that indeed the ancestral *P. maniculatus* was a short-tailed grassland form from which at least two long-tailed forest forms evolved.

We show that the eastern and western forest forms are convergently evolving at the population level. By this, we mean that similar environments (i.e., forests) appear to favor similar phenotypes (i.e., long tails) in two phylogenetic groups that are not closely related to each other (in terms of population differentiation within this species). Furthermore, in two forest-prairie population pairs, one eastern and one western, we find that both pairs differ in the two components of the caudal skeleton that could vary to produce differences in tail length, namely the number of tail vertebrae and the length of those vertebrae. This coupling of vertebral length and vertebral number could be explained in two ways: (1) number and length of vertebrae are controlled by identical, or linked, regions of the genome, or (2) multiple genetic variants controlling number and length of vertebrae have been independently selected in eastern and western populations.

To distinguish between these hypotheses, we examined F2 hybrid individuals from a laboratory intercross between a forest form, *P. m. nubiterrae*, and a prairie form, *P. m. bairdii*. If differences in the length and number of vertebrae are controlled by variants in the same region(s) of the genome, we expect them to be correlated in the F2 individuals. On the other hand, if the two traits are under control of variants in different genomic regions, recombination during the production of gametes in the F1 parents should decouple these traits in the F2 generation, and we should detect no correlation between these traits. We found the latter: we detect no significant correlation between number and length of vertebrae in the F2 (Fig. 5). This result implies that forest environments have favored separate alleles at loci affecting differences in number and length in eastern and western populations. It may be advantageous, biomechanically, to have both more and longer caudal vertebrae when climbing, or it may be simply that longer tails are favored, and alleles affecting length and number were present in the source population from which the forest mice evolved, and thus alleles affecting both traits increased in frequency independently from standing genetic variation in these populations.

There is some precedent for variation in number and length of vertebrae being produced by separate genetic mechanisms. Rutledge *et al*. (1974) performed a selection study in which the authors selected for increased body length and tail length in replicate mouse strains. The authors found that, in two replicate lines selected for increased tail length, one line had evolved a greater number of vertebrae and the other evolved longer vertebrae. That the number and lengths of vertebrae can be genetically uncoupled may not be surprising, given the timing of processes in development that affect these traits. The process of somitogenesis, which creates segments in the embryo that presage the formation of vertebrae, is completed in the *Mus* embryo by 13.5 days of development (Tam 1981), while the formation of long bones does not begin until later in embryogenesis and skeletal growth continues well into the early life of the animal (Theiler 1989).

Together these data convincingly show that longer tails have evolved repeatedly in similar forested habitat, implicating a role of natural selection. Despite separate genetic mechanisms for number and length of vertebrae, both traits contribute approximately equally to the increase in tail length in both the eastern and western forest populations. Future work will explore the biomechanical implications of caudal vertebrae morphology on function (e.g., climbing), and ultimately fitness, as well as the underlying genetic and developmental mechanisms driving this convergently evolved adaptive phenotype.

## Acknowledgements

The authors wish to thank Emily Hager, Jonathan Losos, and Ricardo Mallarino for providing comments on the manuscript. J. Chupasko facilitated work in the MCZ Mammal collection. The following individuals and institutions kindly provided tissue samples (a) and specimen records (b) used in this study: R. Mallarino, L. Turner and A. Young (MCZ; a, b); S. Peurachs (Smithsonian Institution; a, b), C. Dardia (Cornell University Museum of Vertebrates; a, b), S. Hinshaw (University of Michigan Museum of Vertebrates; a, b), L. Olson (University of Alaska Museum of the North; a, b), J. Dunnum (University of New Mexico Museum of Southwestern Biology; a, b), C. Conroy (Museum of Vertebrate Zoology, UC Berkeley; a, b), P. Gegick (New Mexico Museum of Natural History and Science; a, b), S. Woodward (Royal Ontario Museum; a, b), H. Garner (Texas Tech University; a, b), E. Rickart (Utah Museum of Natural History; a, b), R. Jennings (University of Western New Mexico; b), J. Storz (University of Nebraska; a), and the databases of the University of Florida Museum of Natural History (b) and the Burke Museum of Natural History at the University of Washington (b). J. Losos and L. Mahler advised on comparative methods. Read alignment and variant calling were run at the Vital-IT Center (www.vital-it.ch) for high-performance computing of the Swiss Institute of Bioinformatics (SIB). Phylogeographic analyses were run on a server at the School of Life Sciences, University of Cambridge, with the assistance from J. Barna.

## Data archiving

Morphological data and unfiltered VCF, PLINK and TreeMix files are available from the FigShare repository [Reviewer link: figshare.com/s/01f69a425b3911e589ac06ec4b8d1f61]. SNP and mitochondrial alignment files with corresponding phylogenies were deposited in TreeBase (Study 18261) [Reviewer link: http://purl.org/phylo/treebase/phylows/study/TB2:S18261?x-access-code=570d17f6c0a9dee4237ed1023d59441&format=html]">. Mitochondrial sequences were deposited in GenBank (accession numbers KU156828-KU156933).

